# Synthetic lumen rounding directs neural progenitor division mode

**DOI:** 10.64898/2026.03.30.715222

**Authors:** Marina Marchenko, Guillermo Martínez Ara, Juslina Pulikkal, Keisuke Ishihara, Miki Ebisuya

## Abstract

Although function often follows form, the causal role of tissue geometry is difficult to disentangle in complex embryonic development. Brain organoids generated using diverse protocols and species display striking morphological variability, particularly in lumen shape; however, whether and how lumen geometry influences neural development remains unclear. Here, we manipulate lumen sphericity in human cerebral organoids by acutely inducing apical constriction and reveal its impact on the division orientation of apical progenitors. Rapid protein stabilization or optogenetic reconstitution of the apical constriction regulator Shroom3 induces pronounced lumen rounding accompanied by a reduction in apical surface area. In organoids with rounded lumens, apical progenitor divisions shift toward horizontal cleavage planes compared with control organoids, consistent with geometric constraints from the reduced apical surface. Accordingly, rounded-lumen organoids exhibit increased cell delamination and an earlier emergence of basal progenitors in the abventricular region. These findings identify lumen geometry as an instructive regulator of progenitor division mode and lineage progression during early brain development.

## Introduction

Form follows function; yet in developing embryos, function also follows form. Tissue geometry can, for example, sculpt morphogen gradients and modulate mechanosensitive signaling, thereby dictating tissue branching, stem-cell positioning, and differentiation patterns(*1–3*). Disentangling the specific contributions of tissue geometry, however, remains challenging due to multiple interacting processes, confounding factors, and feedback mechanisms inherent to complex developmental systems.

Organoids offer a tractable in vitro platform for manipulating and quantifying tissue geometry. Among them, brain organoids recapitulate key aspects of early neural development, including ventricle-like lumen formation and cell lineage progression(*4–7*). During early cerebral organoid development, neuroepithelial cells transition into radial glial cells; together, they constitute the apical progenitor population that divides at the apical surface(*8–11*). Radial glial cells subsequently undergo asymmetric divisions to generate basal progenitors, including intermediate progenitors, as well as neurons, both of which delaminate from the apical surface and relocate to the abventricular region. Because these asymmetric divisions and delamination events occur at the apical surface, lumen geometry and apical surface area may directly influence cell lineage progression, although the nature and extent of this influence remain unclear.

Consistent with this idea, brain organoids generated using distinct protocols exhibit a broad spectrum of tissue geometries associated with differences in neural development(*12*, *13*). For example, the addition of extracellular matrix (ECM) promotes lumen expansion and telencephalic lineage specification in brain organoids, whereas its absence leads to altered lumen shape and increased neural crest and caudalized identities(*12*). The stem cell source also influences organoid geometry and developmental dynamics: cerebral organoids derived from pluripotent stem cells of microcephaly patients often exhibit reduced overall and lumen size accompanied by premature neurogenesis(*4*, *14*), whereas organoids derived from non-human apes and mice develop more rounded neural buds and lumens as well as faster lineage progression than those derived from humans(*11*, *15–17*). However, variation in organoid protocols and cell sources affects not only tissue geometry but also signaling pathways and cell-fate decisions, thereby confounding causal interpretation.

Determining whether and how tissue geometry instructs neural lineage progression therefore requires strategies that directly and selectively manipulate organoid and lumen geometry while minimizing perturbations to other neurodevelopmental processes. While embedding brain organoids in ECM or hydrogels is commonly used to influence overall organoid geometry, this approach is challenging for directly manipulating lumen architecture, as the lumen is not readily accessible to externally applied materials. An alternative strategy for modulating lumen geometry is apical constriction, an actomyosin-driven contraction of the apical surface of epithelial cells(*18*, *19*). Apical constriction is required for cell shape changes in brain organoids; overexpression of the transcription factor ZEB2 induces apical constriction in neuroepithelial cells, promotes their transition into transitioning neuroepithelial (tNE) cells that subsequently become radial glial cells, and thereby terminates the active proliferative phase(*11*). Because transcription factors exert broad downstream effects, however, a more direct and targeted means of inducing apical constriction is desirable to test its sufficiency and specific roles. Shroom3 is a well-characterized regulator of apical constriction that exerts rapid effects through the recruitment of ROCK to the actomyosin ring at apical junctions(*20–23*). We previously developed an optogenetic tool based on Shroom3, OptoShroom3, to induce apical constriction, which successfully drove lumen shrinkage in mouse optic cup organoids(*24*). We therefore reasoned that Shroom3-based approaches could be leveraged to modulate lumen geometry in human brain organoids.

In this study, we use Shroom3-induced apical constriction in early human cerebral organoids to demonstrate that lumen geometry regulates neural progenitor division mode and the emergence of intermediate progenitors.

## Results

### Shroom3-induced apical constriction drives lumen and bud rounding in cerebral organoids

To examine how sustained apical constriction influences tissue geometry in human cerebral organoids, while avoiding the continuous illumination required for OptoShroom3 activation, we established a chemically inducible Shroom3 system. Shroom3 was fused to a dihydrofolate reductase (DHFR) destabilization domain that is rapidly degraded under basal conditions but stabilized upon addition of the small molecule trimethoprim (TMP)(*25*) (Fig. 1A). Using human induced pluripotent stem cell (iPSC) lines stably expressing Shroom3-DHFR, we generated cerebral organoids following an established protocol, with a reduced initial cell number to generate smaller organoids suitable for high-resolution tissue geometry imaging (fig. S1A). After Matrigel embedding on day 7, protruding neural buds and apical lumens formed around days 8-9, and neurogenesis initiated around days 15-20 (Fig. 1B and fig. S1B). Beginning on day 8, TMP was administered to stabilize Shroom3-DHFR protein (fig. S1, C and D).

**Fig. 1.**
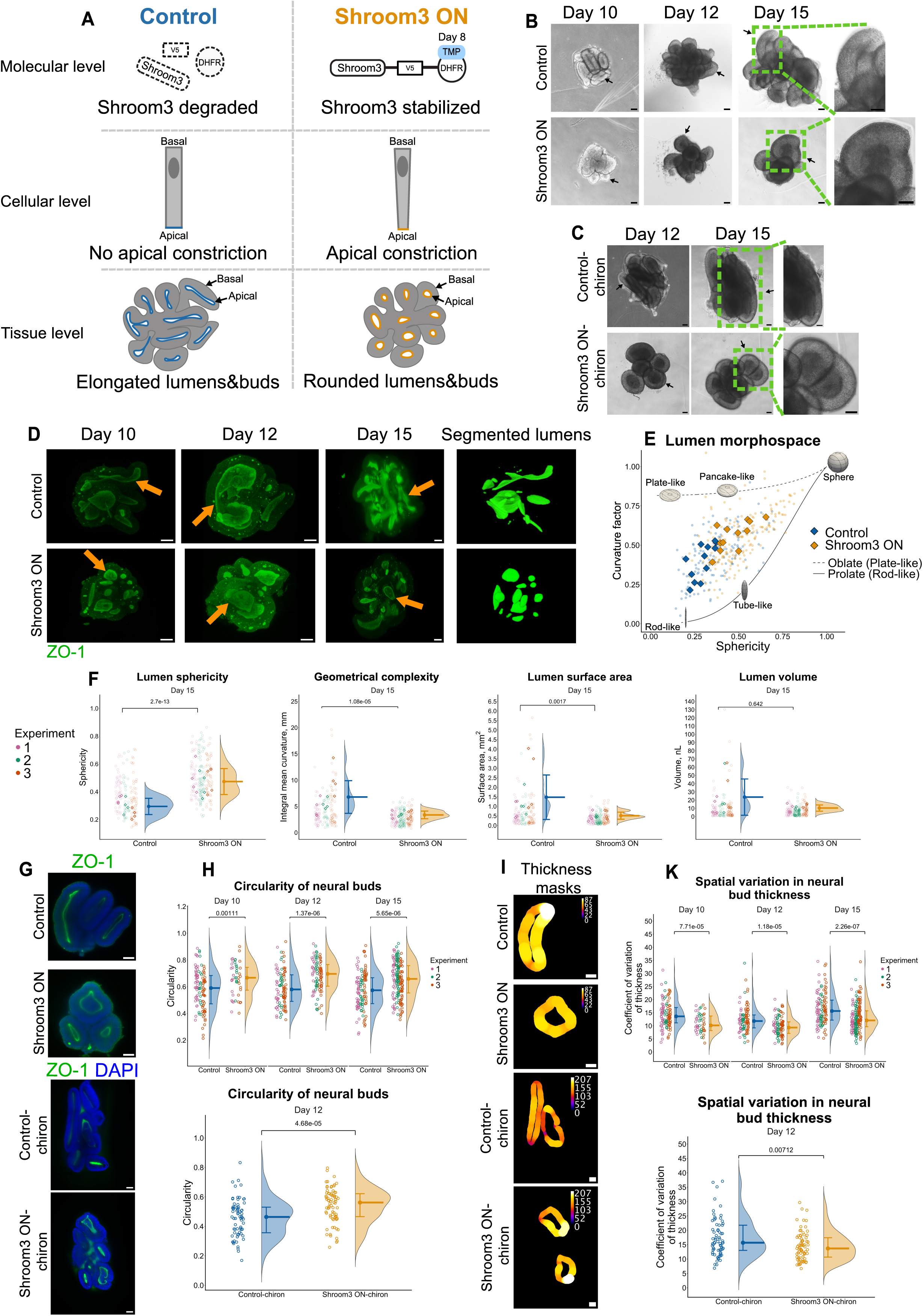
Shroom3-induced apical constriction drives lumen and neural bud rounding in cerebral organoids. (A) Schematic of organoid geometry manipulation. Upon TMP addition, the Shroom3 protein is stabilized, inducing apical constriction. This leads to rounding of lumens and neural buds in cerebral organoids. (B) Representative bright-field images of human cerebral organoid development in the absence (Control) or presence of 10 μM TMP from day 8 (Shroom3 ON). Black arrows indicate differences in neural bud shape between Control and Shroom3 ON conditions. Scale bar: 100 μm. (C) Representative bright-field images of cerebral organoids treated with 3 μM chiron added from day 10. Scale bar: 100 μm. (D) 3D reconstructions of organoids obtained by light-sheet microscopy showing ZO-1 signal (apical marker). Orange arrows indicate lumens. Scale bar: 100 μm. The right panel shows an example of segmented lumens (day 10 sample). Note the difference in scale bars between the day 12 and day 15 images, which results from the use of different microscopes: MuViSPIM (days 10 and 12) and Zeiss light-sheet (day 15). (E, F) Quantification of lumen geometric parameters in day 15 organoids. Raw data are shown as circles; surface area-weighted means as rhombi. Lumens smaller than 0.5 nl were excluded from the analysis. N = 134-186, 12 organoids per condition. (E) Lumen morphospace characterized by lumen sphericity and curvature factor (defined as the ratio of the surface area of a sphere with the same integral mean curvature as the object to the measured surface area, see Methods). (F) Individual lumen geometric parameters. Mean ± SD of the surface area-weighted means is shown. Sphericity: Control = 0.294 ± 0.0585; Shroom3 ON = 0.473 ± 0.0932. Integral mean curvature: Control = 6.78 ± 3.11; Shroom3 ON = 3.37 ± 0.733 mm. Surface area: Control = 1.48 ± 1.17; Shroom3 ON = 0.511 ± 0.186 mm^2^. Volume: Control = 23.6 ± 22.1; Shroom3 ON = 10.3 ± 3.78 nl. (G) Representative immunofluorescence images of neural buds on day 12, in the absence or presence of chiron. (H) Circularity of neural buds. Median and interquartile range (IQR) of circularity: Control day 10 = 0.59 (IQR: 0.472-0.683); Shroom3 ON day 10 = 0.667 (IQR: 0.574-0.745); Control day 12 = 0.579 (IQR: 0.490-0.689); Shroom3 ON day 12 = 0.696 (IQR: 0.604-0.765); Control day 15 = 0.573 (IQR: 0.473-0.667); Shroom3 ON day 15 = 0.658 (IQR: 0.553-0.758). N = 48-172, 9-13 organoids per condition. Control-chiron = 0.464 (IQR: 0.357-0.530); Shroom3 ON-chiron = 0.562 (IQR: 0.466-0.622). N = 69-75, 6 organoids per condition. (I) Thickness of neural buds. Thickness masks were applied to the images shown in panel G. (K) Intra-bud spatial variation in neural bud thickness. Median thickness variation: Control Day 10 = 13.6 (IQR: 11.1-17.0); Shroom3 ON day 10 = 10.1 (IQR: 8.16-13.7); Control day 12 = 11.8 (IQR: 9.18-14.1); Shroom3 ON day 12 = 9.3 (IQR: 7.06-11.5); Control day 15 = 15.6 (IQR: 12.2-19.8); Shroom3 ON day 15 = 12.0 (IQR: 9.63-15.7). Control-chiron = 15.7 (IQR: 13.0-21.8); Shroom3 ON-chiron = 13.7 (IQR: 10.7-17.4). All statistical comparisons were performed using Mann-Whitney U test.

In the absence of TMP (hereafter referred to as the “Control” condition), buds and lumens exhibited a flat and elongated shape (Fig. 1B). In the presence of TMP (the “Shroom3 ON” condition), by contrast, buds and lumens exhibited a more spherical shape (Fig. 1B). To further accentuate these geometric differences, we enhanced apical progenitor proliferation by treating organoids with the WNT signaling agonist chiron(*26*, *27*). Chiron treatment generally increased organoid size; however, Control organoids formed buds and lumens with an even more flattened and elongated shape, while Shroom3 ON buds maintained a spherical shape (Fig. 1C).

We quantified three-dimensional (3D) lumen geometry from light-sheet microscopy images of ZO-1-stained samples using a machine learning-based segmentation pipeline and previously described geometric parameter estimation(*28*) (fig. S2A). Control organoids often displayed interconnected, flat, and elongated lumens, whereas Shroom3 ON organoids formed separated, geometrically simpler, and more spherical lumens (Fig. 1, D and E, and movie S1). These geometric differences first became evident on day 10, two days after the induction of apical constriction, and persisted until at least day 15. Quantitatively, Shroom3 ON lumens exhibited 1.6-fold higher sphericity and 2-fold lower integral mean curvature, a metric of surface geometric complexity, compared with Control lumens (Fig. 1F and fig. S2B). Consistent with these observations, lumen surface area in Shroom3 ON organoids was 2.9-fold smaller than that in Control organoids, whereas lumen volume did not differ significantly between conditions (Fig. 1F and fig. S2B).

Consistent with the differences in lumen geometry, 2D cross-sectional analysis showed that neural buds in Shroom3 ON organoids had higher circularity than Control buds, both in the absence and presence of chiron (Fig. 1, G and H). Mean bud thickness was higher in Shroom3 ON buds on day 10 but became comparable between conditions by day 12 (fig. S2C). Notably, Shroom3 ON buds showed reduced intra-bud spatial variability in thickness, a feature of a more homogeneous and geometrically simpler structure (Fig. 1, I and K). These results indicate that Shroom3 ON buds adopt a more rounded geometry than Control buds. At the scale of the entire organoid, Shroom3 ON organoids tended to exhibit increased sphericity and reduced total volume; however, unlike the pronounced changes observed in lumen and bud geometry, these organoid-level differences were modest and not consistently significant (fig. S2D). The rounding of lumens and buds was independently reproduced in organoids derived from a second iPSC clone expressing Shroom3-DHFR (fig. S2E). TMP treatment alone, in the absence of Shroom3-DHFR, did not alter lumen and bud geometry, confirming that the observed effects were attributable to Shroom3 stabilization rather than to TMP itself (fig. S2F).

Together, these results demonstrate that Shroom3-induced apical constriction is sufficient to drive the formation of rounded, geometrically simplified lumens and neural buds in human cerebral organoids.

### Tissue rounding accelerates basal progenitor generation

We next assessed how altered tissue geometry impacts progenitor cell division and differentiation patterns. At this stage, most mitotic cells are radial glial cells, an apical progenitor subtype that divides at the apical surface. However, we occasionally detected phospho-histone H3 (PH3)-positive mitotic cells in the abventricular region of day 15 Shroom3 ON organoids (Fig. 2A). Notably, Shroom3 ON organoids exhibited a 2.1-fold increase in abventricular dividing cells compared with Control organoids (Fig. 2B). Among these abventricular dividing cells, 85.3% were SOX2-positive and TBR2-negative, suggesting that they originate from apical progenitors (radial glial or neuroepithelial cells) that have delaminated from the apical surface (Fig. 2C). By contrast, 14.7% of abventricular dividing cells were SOX2-negative and TBR2-positive, indicative of intermediate progenitor identity (Fig. 2C). This increase in abventricular cell divisions under the Shroom3 ON condition was also observed in chiron-treated organoids (Fig. 2, D and E). These results suggest a link between apical surface reduction and increased progenitor relocation to the abventricular region in Shroom3 ON organoids. By contrast, the cleaved CASPASE-3 was detected in only a small fraction of cells, suggesting no evident difference in apoptosis between conditions (fig. S3A).

**Fig. 2.**
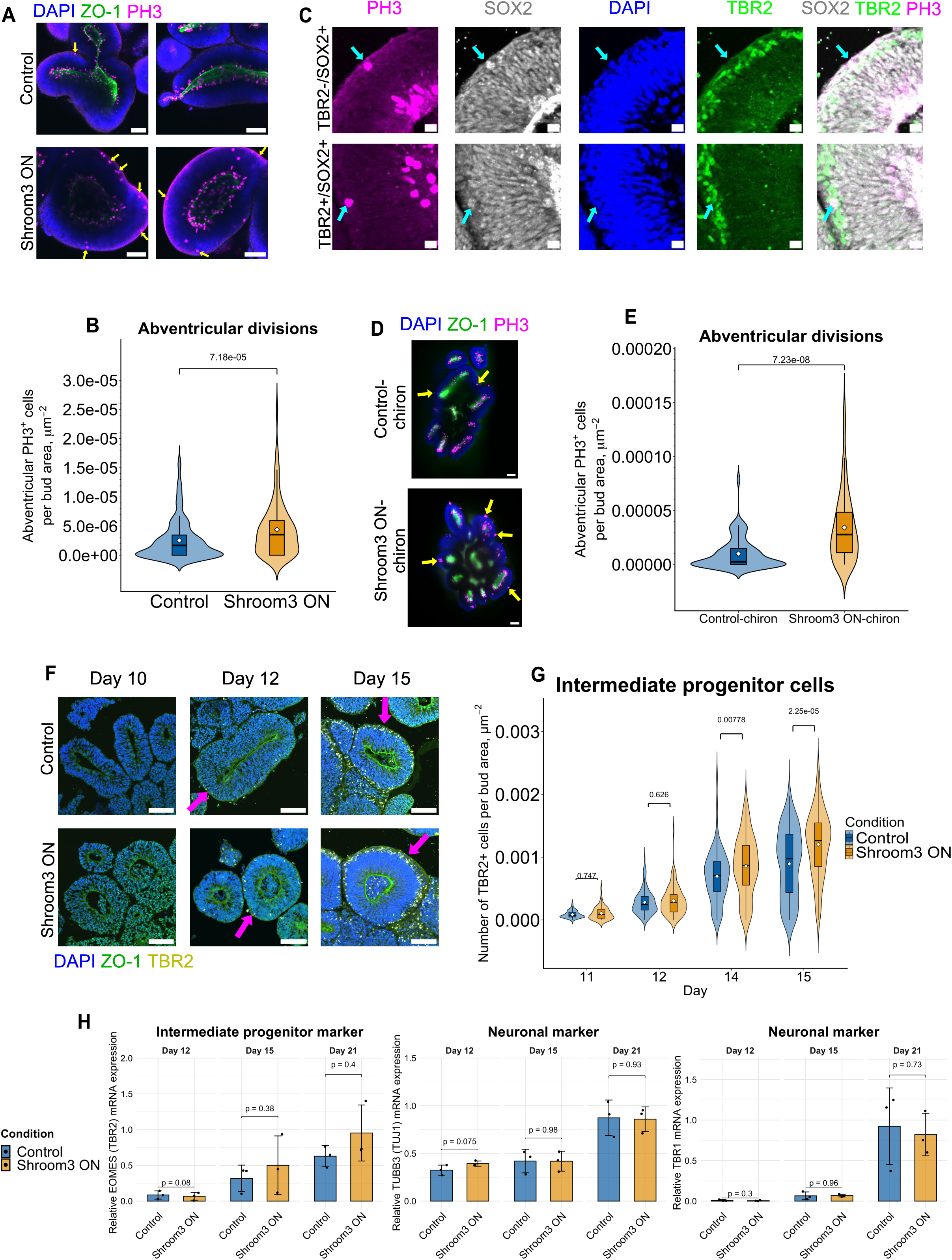
Tissue rounding accelerates basal progenitor generation. (A) Representative immunofluorescence images of day 15 neural buds, labeled for PH3 (mitotic cells, magenta), DAPI (nuclei, blue), and ZO-1 (apical marker, green). Yellow arrows indicate abventricular cell divisions. Scale bar: 100 μm. (B) Quantification of PH3-positive cells in the abventricular region normalized to bud area. Median abventricular divisions: Control = 0.00000168 (IQR: 0-0.00000346); Shroom3 ON = 0.00000354 (IQR: 0-0.00000593). N = 115-172, 12-13 organoids per condition. (C) Representative immunofluorescence images of day 15 neural bud cryosections. Among the abventricular PH3-positive cells, TBR2-negative and SOX2-positive cells represent putative apical progenitors delaminated from the apical surface (upper panel). TBR2-positive and SOX2-positive cells correspond to intermediate progenitor cells (lower panel). Scale bar: 15 μm. (D) Representative immunofluorescence images of day 12 organoids treated with chiron. Yellow arrows indicate abventricular cell divisions. Scale bar: 100 μm. (E) Median abventricular divisions: Control-chiron = 0.0000026 (IQR: 0-0.0000149); Shroom3 ON-chiron = 0.0000278 (IQR: 0.000011-0.0000484). N = 68-75, 6 organoids per condition. (F) Representative immunofluorescence images of neural bud cryosections, labeled for TBR2 (intermediate progenitors, yellow), DAPI (nuclei, blue), and ZO-1 (apical marker, green). Magenta arrows indicate TBR2-positive cells. Scale bar: 100 μm. (G) Quantification of TBR2-positive cells normalized to bud area from day 11 to day 15. N = 50-140, 11-31 organoids per condition. Median number of TBR2-positive cells: Control Day 11 = 0.0000675 (IQR: 0.0000459-0.000124); Shroom3 ON day 11 = 0.0000834 (IQR: 0.0000264-0.000170); Control day 12 = 0.000245 (IQR: 0.000158-0.000376); Shroom3 ON day 12 = 0.000282 (IQR: 0.000128-0.000403); Control day 14 = 0.000708(IQR: 0.000453-0.000930); Shroom3 ON day 14 = 0.000871 (IQR: 0.000554-0.00119); Control day 15 = 0.000974(IQR: 0.000437-0.00137); Shroom3 ON day 15 = 0.00126 (IQR: 0.000853-0.00155). (B, E, G) Statistical comparisons were performed using the Mann-Whitney U test. (H) Quantitative RT-PCR analysis of TBR2, TUBB3, and TBR1 mRNA levels in Control and Shroom3 ON organoids. Mean ± SD. N = 3, 30 organoids were pooled per condition. Statistical comparisons were performed using the two-sided paired t-test.

Motivated by the increase in abventricular divisions, we quantified intermediate progenitors, the earliest basal progenitor subtype. TBR2-positive intermediate progenitors first appeared around day 12 in Shroom3 ON organoids, whereas they were barely detectable in Control organoids (Fig. 2F). On days 14 and 15, Shroom3 ON organoids consistently exhibited 1.2-1.3-fold higher numbers of intermediate progenitors than Control organoids (Fig. 2G). The increase in intermediate progenitors in Shroom3 ON organoids was reproduced independently using the second Shroom3-DHFR iPSC clone (fig. S3B). TMP treatment alone, in the absence of Shroom3-DHFR, did not significantly alter the number of intermediate progenitors (fig. S3C).

To determine whether the accelerated lineage progression in Shroom3 ON organoids persisted at later stages, we quantified the expression of neuronal markers. The intermediate progenitor marker TBR2 showed a trend toward higher expression in Shroom3 ON organoids compared with Control organoids, although this difference did not reach statistical significance (Fig. 2H). By contrast, the neuronal markers TUBB3 and TBR1 showed no significant differences between conditions (Fig. 2H). The absence of detectable differences in neuronal marker expression may reflect the limited sensitivity of bulk qRT-PCR or, alternatively, compensatory neurogenesis in Control organoids potentially driven by the onset of endogenous SHROOM3 expression around day 15 in human cerebral organoids(*11*).

Together, these results reveal a strong association between more spherical tissue geometry and accelerated basal progenitor generation.

### Rounded lumen geometry shifts apical progenitor divisions toward horizontal cleavage planes

We next investigated the mechanisms by which lumen and bud rounding promote basal progenitor generation. Because intermediate progenitors arise from asymmetric divisions of radial glial cells at the apical surface, we examined their division modes. Cleavage plane orientation at anaphase is a key determinant of daughter cell fate: vertically oriented cleavage planes (near 90° relative to the apical surface) typically result in symmetric divisions that produce two radial glial cells(*8*, *29–33*) (Fig. 3A, Upper). By contrast, oblique (near 45°) or horizontal (near 0°) cleavage planes are strongly associated with asymmetric division, in which one daughter cell retains radial glial identity while the other exits the ventricular zone to become an intermediate progenitor or a neuron(*8*, *29–33*) (Fig. 3A, Lower).

**Fig. 3.**
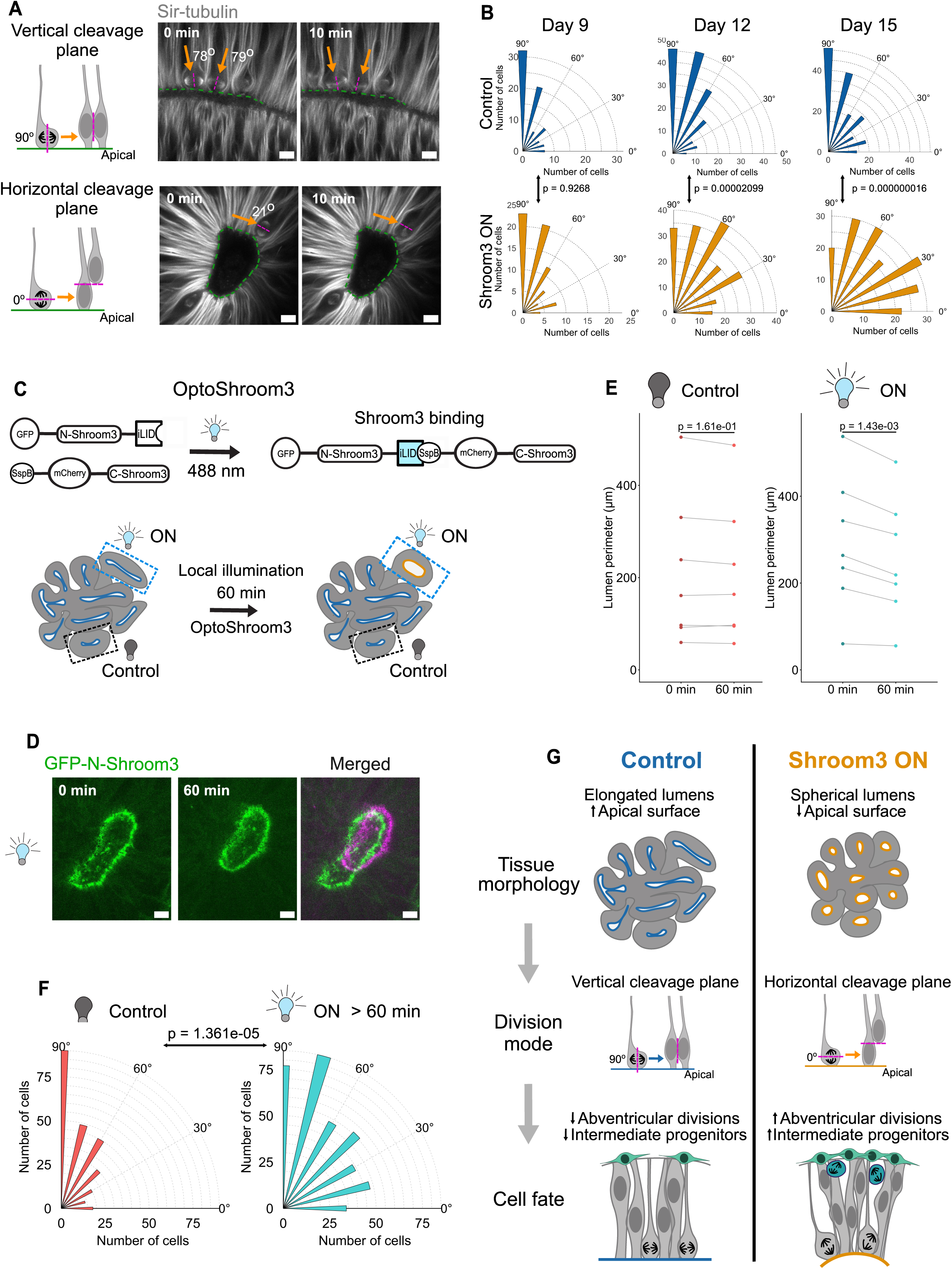
Rounded lumen geometry shifts apical progenitor divisions toward horizontal cleavage planes. (A) Schematic of apical progenitor division mode (left). Purple dashed lines indicate the cleavage planes. Vertical cleavage planes keep both daughter cells attached to the apical surface (upper panel), whereas horizontal cleavage planes lead to the delamination of one daughter cell (lower panel). Representative live-imaging frames of day 12 cerebral organoids used for cleavage orientation measurements (right). Mitotic spindles were visualized with SiR-tubulin. Green dashed lines indicate the lumen surface, and orange arrows track dividing cells across time points. Scale bar: 10 μm. (B) Distribution of cleavage plane orientations in Control and Shroom3 ON conditions. Measurements were taken at the anaphase, when microtubules start to separate. Angles near 90° correspond to vertical cleavage planes, whereas angles near 0° correspond to horizontal cleavage planes. Angles were grouped into seven categories: 0°, 15°, 30°, 45°, 60°, 75°, and 90°. N = 79-195, 9-12 organoids per condition per day. Watson-Williams test was performed to compare conditions. (C) Schematic of the OptoShroom3 experiment. Constructs containing the N-terminus (GFP-N-Shroom-iLID) and C-terminus (SspB-mCherry-C-Shroom3) of the Shroom3 protein bind to each other upon blue-light illumination (upper panel). The reconstituted full-length Shroom3 induces apical constriction and lumen rounding in illuminated buds (ON), but not in non-illuminated buds (Control) within the same organoid (lower panel). (D) Representative imaging frames of illuminated lumens in a day 12 organoid visualized with GFP-N-Shroom3-iLID, which constitutively localizes to the apical surface. Light stimulation was applied in 9-min ON / 1-min OFF cycles. The left and center images show the start and end of 60 min of photostimulation, respectively, and the right image shows the overlay of the two frames. Scale bar: 10 μm. (E) Lumen perimeter measurements in day 12-15 organoids before and after OptoShroom3 photostimulation. The light bulb icons indicate non-illuminated (Control, dark) and illuminated (ON, blue) lumens within the same organoid. N = 7. Two-sided paired t-test was performed to compare perimeters before and after photostimulation. (F) Distribution of cleavage plane orientations in Control and ON neural buds in day 12-15 organoids, measured as in panel B. The measurements started after 60 min of OptoShroom3 activation. N = 266-397, 7 neural buds per condition. Watson-Williams Test was performed to compare conditions. (G) Schematic of the role of lumen rounding in regulating cell division mode and cell fate. In the bottom panel, green, turquoise, and gray cells represent intermediate progenitors (basal progenitors), abventricular dividing cells, and radial glial cells (apical progenitors), respectively.

On day 9, apical progenitors in both Control and Shroom3 ON organoids predominantly divided with vertically oriented cleavage planes (Fig. 3B). By day 12, however, cleavage plane orientation shifted toward oblique and horizontal angles, with this shift being more pronounced in Shroom3 ON organoids (Fig. 3B, fig. S4, and movie S2). This enhanced bias toward horizontal cleavage planes in Shroom3 ON organoids remained evident on day 15 (Fig. 3B). The timing of this shift in division mode coincided with the emergence of abventricular divisions and intermediate progenitors in Shroom3 ON organoids (Fig. 2G), supporting the idea that radial glial cells with horizontally oriented cleavage planes frequently undergo asymmetric divisions to generate intermediate progenitors.

To independently test the link between lumen geometry and apical progenitor division mode, we employed an alternative approach. Because stabilization of Shroom3-DHFR requires TMP and may introduce side effects associated with Shroom3 overexpression, we used OptoShroom3, which relies on light-induced interaction between the N- and C-terminal Shroom3 fragments mediated by iLID-SspB binding(*24*) (Fig. 3C). The N-Shroom3-iLID fragment constitutively localizes to the apical surface, whereas the SspB-C-Shroom3 fragment rapidly translocates upon light illumination, reconstituting full-length Shroom3 and inducing apical constriction within minutes, without altering overall Shroom3 expression levels(*24*). After 60 min of localized light illumination of neural buds, OptoShroom3 organoids exhibited more rounded lumens and an average 13% reduction in lumen perimeter, consistent with results obtained using Shroom3-DHFR (Fig. 3, D and E, fig. S5, and movie S3). Notably, illuminated OptoShroom3 buds showed a pronounced shift in apical progenitor divisions toward oblique and horizontally oriented cleavage planes compared with non-illuminated buds within the same organoids (Fig. 3F and movie S4). Given the short time course (within 5 h of illumination onset), we conclude that OptoShroom3 activation directly modulates division mode rather than acting through long-term signaling or gene expression-dependent processes.

Together, these results suggest that lumen rounding and apical surface shrinkage constrain divisions with vertically oriented cleavage planes, thereby promoting a shift toward horizontally oriented cleavage planes (Fig. 3G). This shift in division orientation of apical progenitors likely promotes their delamination from the apical surface, increasing abventricular divisions and accelerating the generation of basal progenitors, including intermediate progenitors.

## Discussion

In this study, we show that apical constriction reshapes the lumen geometry of human cerebral organoids and that this change biases the division mode of apical progenitors, thereby accelerating lineage progression toward basal progenitors (Fig. 3G).

While previous studies have shown that lumen expansion in brain organoids is accompanied by activation of WNT/YAP signaling(*12*) and that apical constriction is required for neuroepithelial cell shape changes(*11*), it has remained unclear whether and how changes in lumen geometry driven by apical constriction influence cell fates during early brain development. Here, we find that lumen rounding and the accompanying reduction in apical surface area bias apical progenitor divisions toward horizontal cleavage planes. This bias may arise from reduced available space for vertically oriented divisions or, alternatively, from alterations in the apical cortex that affect spindle positioning. Although the precise molecular mechanisms underlying this bias require further investigation, our work provides a conceptual framework for understanding how geometric constraints influence cell fate decisions.

The observed shift in division orientation coincided with an increase in basal progenitors, suggesting that lumen rounding and apical surface shrinkage accelerate the transition from apical to basal progenitors. Brain organoids derived from microcephaly patients, as well as those derived from non-human apes and mice, are characterized by smaller apical surface areas and exhibit faster lineage progression and earlier shifts in division mode(*4*, *11*, *14*, *16*, *17*), raising the possibility that similar geometric mechanisms operate across systems. Despite the increase in intermediate progenitors, we did not detect clear downstream effects of lumen rounding on neuronal output at later stages. Nevertheless, Shroom3 has previously been shown to promote neurogenesis in the zebrafish retina(*34*). The absence of a clear neuronal increase in Shroom3 ON organoids may therefore reflect gradual compensatory neurogenesis in Control organoids or limited sensitivity of bulk qRT-PCR to detect subtle changes in cell type composition. Higher-resolution approaches, such as single-cell RNA sequencing, will be required to assess whether lumen geometry affects long-term neuronal output.

We manipulated lumen and bud geometry by selectively inducing apical constriction through Shroom3, providing a genetic approach to control tissue geometry. In the emerging field of synthetic morphogenesis, direct manipulation of tissue architecture remains technically challenging, and genetic tools for precise regulation of cell and tissue shape are still limited(*35–38*). Although optogenetic recruitment of RhoGEFs to the apical membrane is well established for inducing apical constriction in Drosophila embryos(*39*), selective illumination of the apical membrane is difficult in brain tissues with inward-facing apical surfaces. Shroom3 is particularly well suited for this purpose, as it localizes only to apical junctions and induces apical constriction without acting as a transcriptional regulator, thereby minimizing broad perturbations to gene expression programs. OptoShroom3 enabled rapid, spatially confined manipulation of lumen geometry, whereas stabilization of Shroom3-DHFR produced sustained geometric changes and altered lineage progression. The organoid system further provided experimental advantages, as organoids are highly amenable to external manipulation. Despite inherent batch-to-batch variability, our findings were validated across multiple organoid batches and two independent cell clones. Moreover, OptoShroom3 enabled direct comparisons between illuminated and non-illuminated buds within the same organoid, providing an internal control.

Shroom3-induced apical constriction may affect processes beyond tissue geometry, including mechanical tension and downstream signaling pathways. However, the rapid onset of division orientation changes following OptoShroom3 activation argues against slower, transcription-dependent signaling mechanisms. Shroom3-induced apical constriction may also increase basal progenitors by accelerating the transition from neuroepithelial cells to tNE cells, which subsequently give rise to radial glial cells(*11*). Nevertheless, the overall increase in abventricular dividing cells, including SOX2-positive and TBR2-negative putative delaminated apical progenitors, suggests that intermediate progenitor expansion primarily results from enhanced cell delamination and asymmetric divisions, rather than from an accelerated transition to tNE cells.

Future challenges include developing strategies to manipulate lumen geometry independently of mechanical tension, thereby disentangling the respective contributions of geometry and force to division mode. Another important direction will be to investigate whether lumen geometry similarly influences lineage progression in the developing brain in vivo. Overall, our work establishes lumen geometry as an instructive and engineerable regulator of progenitor division modes and lineage progression during neural development, rather than a passive byproduct of morphogenesis. We anticipate that similar geometric principles operate during later stages of brain development and across diverse morphogenetic processes throughout the body.

## Methods

### Human induced pluripotent stem cell (iPSC) culture

Human iPSC line (iPS(IMR90)-4) was obtained from WiCell. iPSCs were maintained in StemFlex medium (Fisher Scientific, A3349401) on Matrigel-coated dishes in a humidified incubator at 37 °C with 5% CO_2_. DMEM-F12 (Gibco/Fisher Scientific, 11330-032/11574546) with 1.23% Matrigel (Corning, 356231) was used to coat plates for 30 min at 37 °C prior to cell passaging. Cells were passaged every 3-4 days with Accutase (Gibco/Fisher Scientific, A1110501/11599686) at a density of 200,000 cells per well in a 6-well plate (Corning, 353046). The StemFlex medium was supplemented with ROCK inhibitor Y-27632 (10 μM) (StemCell Technologies, 72308) during the first day after passaging to enhance cell survival. The culture medium was changed daily, with 2 ml per well. Cells were regularly tested for mycoplasma contamination and cryopreserved in Stem-CellBanker (Amsbio, 11890) in liquid nitrogen for long-term storage.

All experiments involving human iPSCs were approved by the University Hospital Ethical Review Committee (IRB00001473; IORG0001076; approval number SR-EK-102022025).

### Generation of genetic constructs and clonal cell lines

The mouse Shroom3 sequence was a gift from T. Nishimura from M. Takeichi lab. The DHFR sequence was obtained from the ProteoTuner Guard system (Clontech)(*25*). The Shroom3-V5-DHFR construct was generated using the MultisiteGateway system (ThermoFisher) and the PiggyBac destination vector(*40*) with the CAG promoter.

The Shroom3-V5-DHFR sequence is shown below.

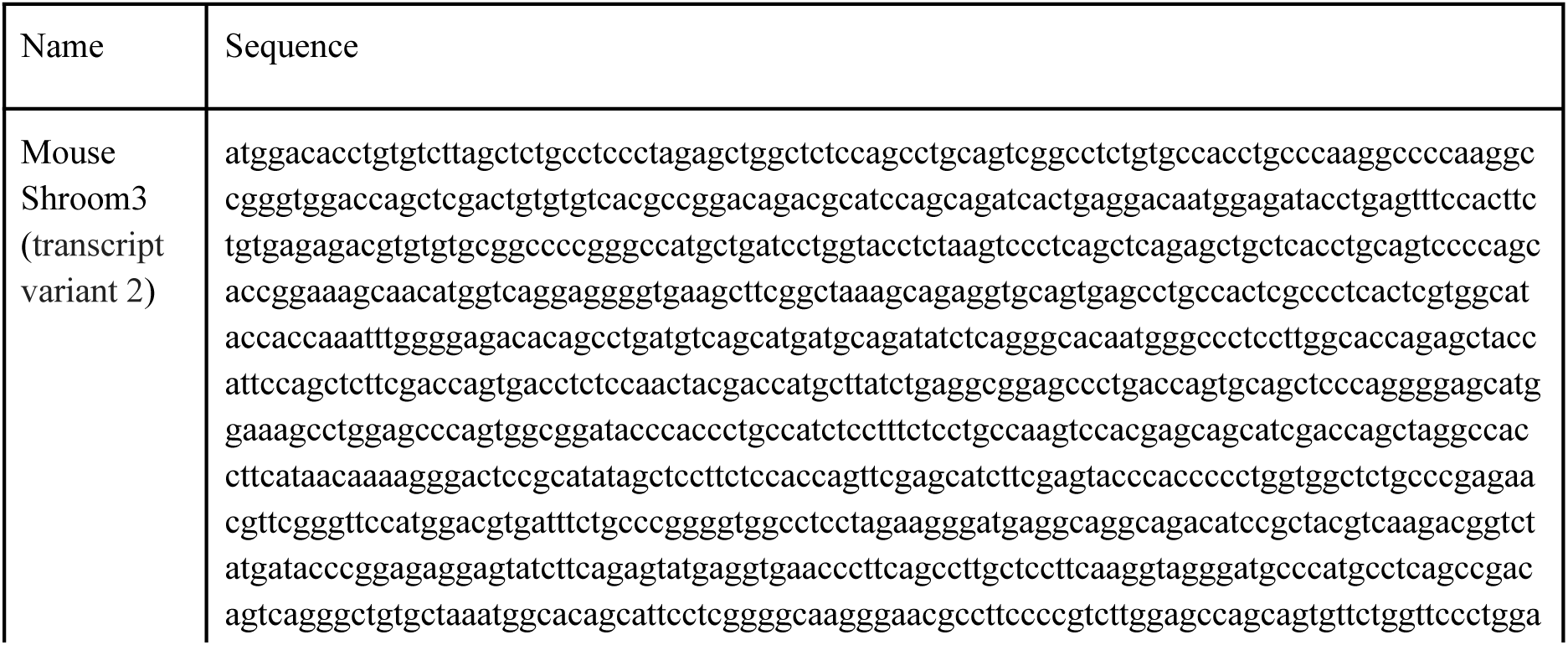
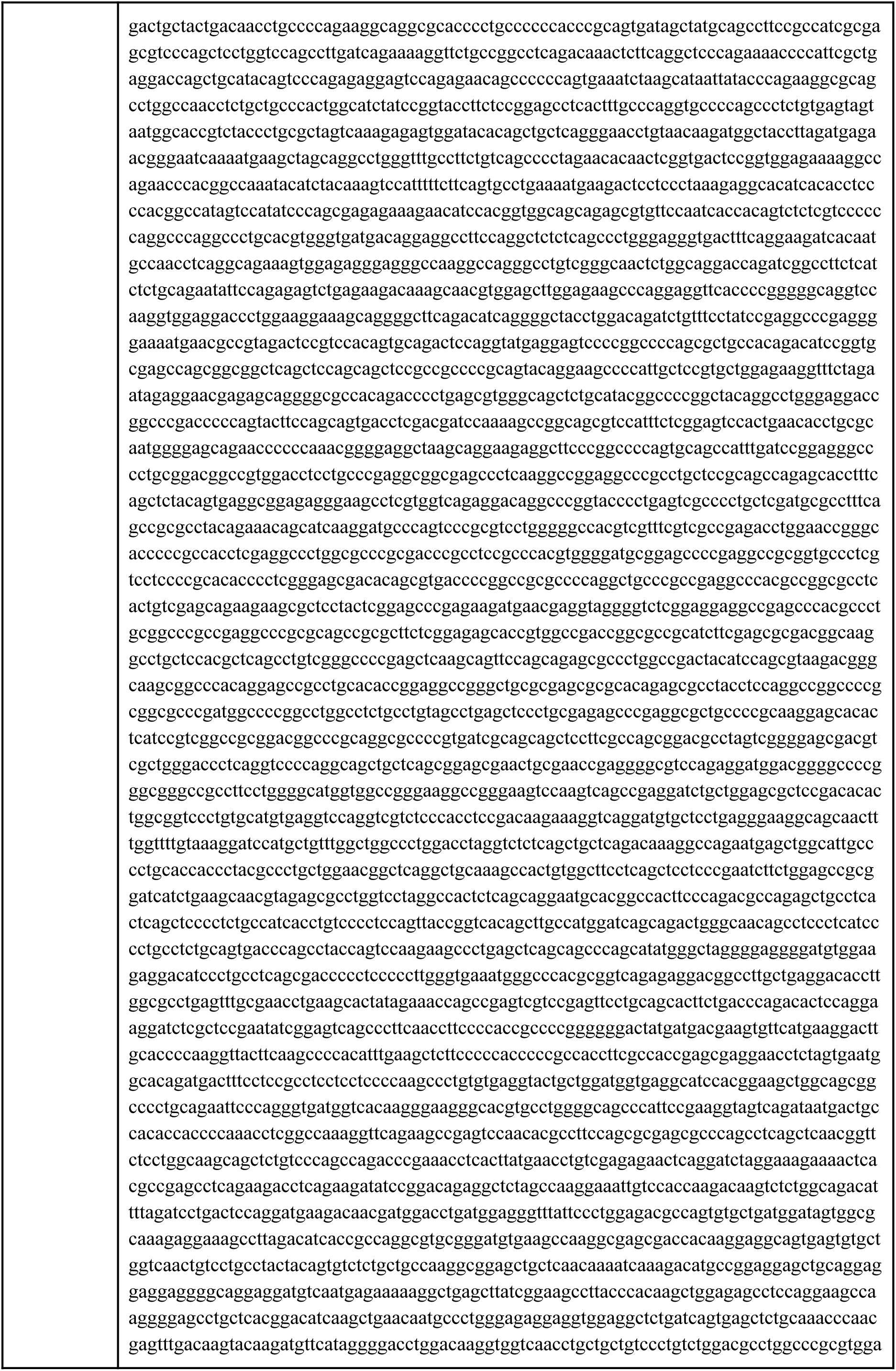
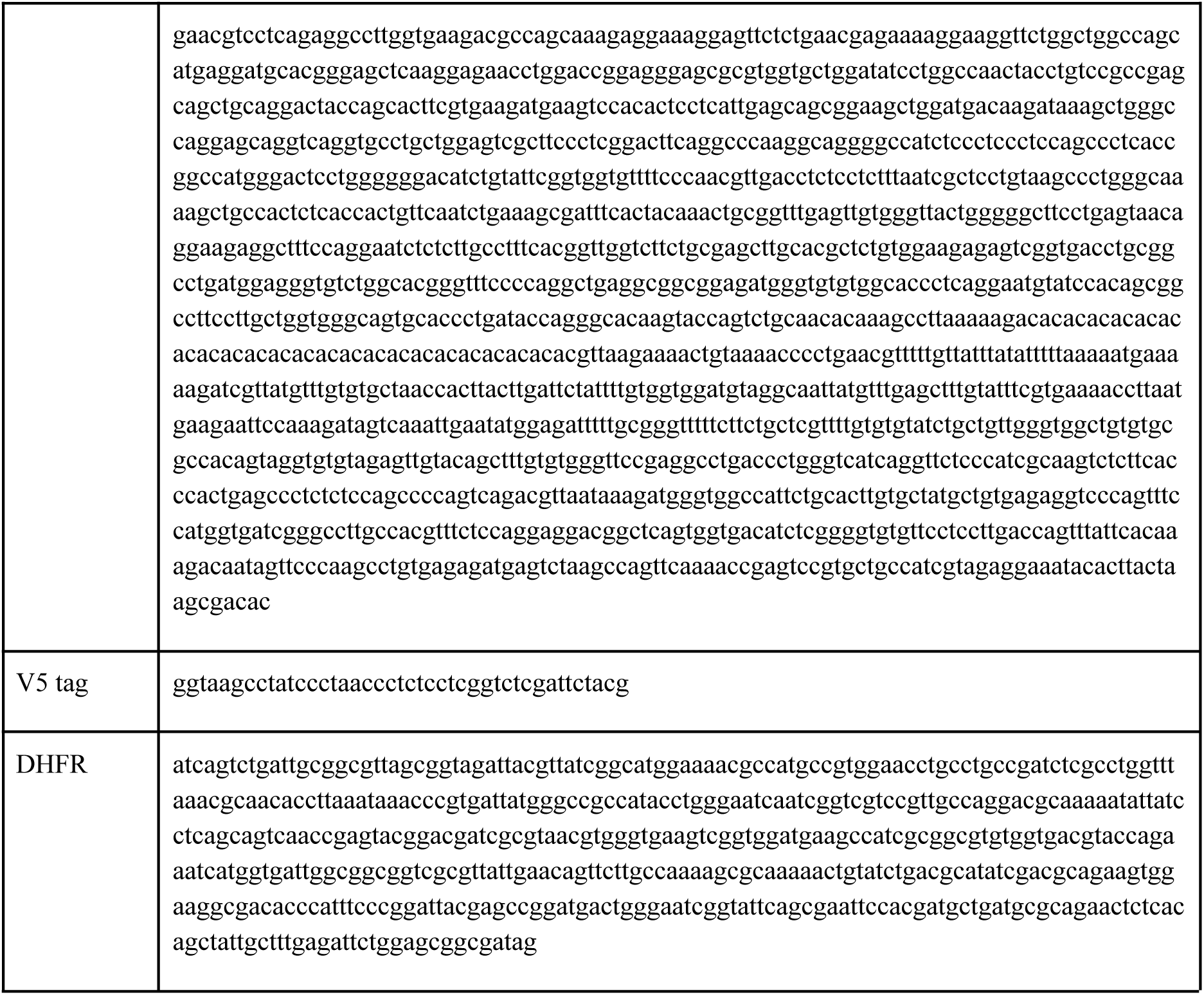

For the generation of the Shroom3-DHFR iPSC line, the construct was transfected using the PiggyBac system and Amaxa 4D Nucleofector (Lonza). A total of 500,000 iPSCs were resuspended in 20 μL P3 Primary Cell Solution (P3 Primary Cell 4D-Nucleofector X Kit, Lonza), mixed with 0.5 μg of Shroom3-V5-DHFR plasmid and 0.5 μg of PiggyBac transposase plasmid, and electroporated. After electroporation, cells were plated on Matrigel-coated dishes and cultured in the StemFlex medium. To obtain clonal lines, cells were transferred to laminin-coated dishes (Amsbio, AMS.892-021) at a low density (200-400 cells per 6 cm dish) in the medium supplemented with ROCK inhibitor. When colonies developed well-defined edges and morphology, they were manually picked under a bright-field microscope and transferred to 96-well plates. Colonies were screened for uniformly high Shroom3 expression by immunostaining. The OptoShroom3 iPSC line was previously described(*24*), and the corresponding GFP-NShroom3-iLID and SspB-mCherry-CShroom3 constructs are available from Addgene (plasmid #170976 and #170977).

### Cerebral organoid generation

Cerebral organoids were generated from human iPSC lines using the STEMdiff Cerebral Organoid Kit (StemCell Technologies, 08570). To generate smaller organoids, the initial cell number was reduced to 150-250 cells. The original protocol (StemCell Technologies Document #DX21849) was followed with minor modifications; on the day of embryoid body formation (day 0), cells were centrifuged for 5 min at 300 × g to ensure better cell aggregation (fig. S1A). To enhance cell proliferation, 3 μM chiron (CHIR99021, TOCRIS, 4423) was added on day 10.

### Manipulation of organoid geometry

To generate cerebral organoids with controlled geometry, two Shroom3-DHFR clonal iPSC lines were used, with clone 1 used for most experiments and clone 2 used for validation. Following Matrigel embedding, 10 μM trimethoprim (TMP; ThermoFisher, 455120050) was added on day 8 to stabilize the Shroom3 protein.

To activate apical constriction with OptoShroom3, an Olympus FV3000 confocal microscope was used. Local bud illumination was performed at 0.1% power with a 488-nm laser in all samples. UPLSAPO10X2 and UPLSAPO30XS (silicon oil) objectives were used for organoid imaging and photostimulation.

### Immunofluorescent staining of human iPSCs

Shroom3 protein stabilization in clonal cell lines was evaluated by immunostaining of Shroom3-DHFR iPSC lines (fig. S1C). Cells were cultured on imaging-compatible plastic dishes (Ibidi, 81816) in the presence of TMP for 24 h and subsequently fixed in 3.7% formaldehyde (Sigma-Aldrich, 1040031000) for 10 min at room temperature. Samples were washed three times with phosphate-buffered saline (PBS) and permeabilized with 0.5% Triton X-100 (Sigma-Aldrich, T9284-100ml) for 10 min. After washing with PBS, samples were incubated with the anti-V5 tag antibody (1:300) in 3% BSA solution (Proliant, 68100-PHB) for 1 h at room temperature, followed by incubation with the secondary antibody (1:300) for 1 h at room temperature. Samples were washed three times with PBS and imaged using the Olympus FV3000 confocal microscope with UPLSAPO10X2 or UPLSAPO30XS (silicon oil) objectives.

### Immunofluorescent staining of organoid cryosections

Cerebral organoids were fixed in 4% paraformaldehyde (PFA) (ThermoFisher, 28908) either at room temperature for 20 min or overnight at 4 °C. Samples were then washed three times for 5 min each with PBS. For cryostat processing, samples were incubated overnight in 30% sucrose (Sigma-Aldrich, S0389-500G) in PBS, embedded in OCT compound (Weckert, 600001), frozen at -20 °C overnight, and cryosectioned at a thickness of 14 μm using a CryoStar NX70 cryostat. The sectioned samples were collected onto glass slides (R. Langenbrinck GmbH, 03-0060) and stored at -20 °C.

Cryosections were thawed for 30 min at room temperature in a humidified chamber. Samples were then washed three times with PBS. The areas containing the sections were marked using a hydrophobic marker (BIOZOL, VEC-H-4000) to prevent sample drying during the staining procedure. Blocking and permeabilization were performed by adding TX blocking solution for 15 min at room temperature. The primary antibody diluted in blocking solution (1:200) was added to the slides overnight at room temperature. After washing three times with PBS, the secondary antibody (1:200) was added and incubated for at least 2 h at room temperature. The slides were washed with PBS and mounted with coverslips (Marienfeld Superior, 101242), then sealed with transparent nail polish to prevent sample drying.

Cryosections were imaged using the Olympus FV3000 confocal microscope with UPLSAPO10X2 or UPLSAPO30XS (silicon oil) objectives.

### Immunofluorescent staining and imaging of whole-mount organoids

Cerebral organoids were fixed in 4% PFA either at room temperature for 20 min or overnight at 4 °C. They were then washed with PBS three times for 5 min. Blocking and permeabilization were performed using TX blocking solution (0.2% Triton X-100, 4% normal donkey serum (abcam, ab7475) in PBS) overnight at 4 °C in a 1.5 ml Eppendorf tube. Organoids were then incubated with the primary antibody diluted in blocking solution (1:200) overnight at 4 °C. After washing with PBS three times for 5 min, organoids were incubated with the secondary antibody (1:200) diluted in TX blocking solution overnight at 4 °C.

For imaging with the Olympus FV3000 confocal microscope, the organoids were transferred to imaging plates (Sigma, CLS4580-10EA) and imaged with UPLSAPO10X2 or UPLSAPO30XS (silicon oil) objectives.

For imaging with the Luxendo MuViSPIM microscope, the organoids were embedded in 2% low-melting-point agarose (Sigma-Aldrich, A9414-5G) and mounted within FEP tubes (Karl Schupp AG, inner diameter 1.7 mm, wall thickness 0.05 mm). 3D imaging was performed with two Nikon Plan Apo 4x illumination objectives and two CFI90 20xC Glyc detection objectives.

For imaging with the Zeiss LightSheet microscope, the organoids were incubated overnight in a clearing solution(*41*) with a refractive index (RI) of 1.48 at room temperature. To prepare a clearing solution, 33 ml glycerol, 7 ml distilled water, and 29.72 g sucrose, instead of fructose, were mixed using a magnetic stirrer until fully dissolved. The clearing solution was equilibrated to room temperature before use in the imaging chamber. This clearing protocol results in minimal changes to organoid size. Samples were embedded in 2% low-melting-point agarose and mounted in clearing solution within glass capillaries (green, BRAND). Illumination was achieved with two Zeiss 10x air objectives (NA 0.2) with refractive index correction ranging from 1.33 to 1.58. Imaging was performed using a 20x clearing immersion objective (NA 1.0, RI = 1.48 ± 0.03; working distance 5.6 mm).

### Antibodies used in this study

#### Primary antibodies

- V5 Tag Monoclonal Mouse Antibody (SV5-Pk1) (ThermoFisher, R960-25)

- Monoclonal Mouse Human/Mouse/Rat SOX2 antibody (R&D systems, MAB2018)

- Anti-Histone H3 (phospho S28) Rat antibody [HTA28] (abcam, ab10543)

- Anti-TBR2/Eomes antibody [EPR21950-241] (abcam, ab216870)

- Recombinant Anti-PAX6 antibody [EPR15858] (abcam, ab195045)

- anti-Tubulin β3 (TUBB3) Antibody (Previously Covance, MMS-435P)

- ZO-1 Monoclonal Mouse Antibody (ZO1-1A12), Alexa Fluor 647 (ThermoFisher, MA3-39100-A647)

- Cleaved Caspase-3 (Asp175) (5A1E) Rabbit mAb (Cell Signaling Technology, #9664)

#### Secondary antibodies

- Goat anti-Mouse IgG (H+L) Highly Cross-Adsorbed Secondary Antibody, Alexa Fluor 488 (ThermoFisher, A-11029)

- Goat anti-Mouse IgG (H+L) Highly Cross-Adsorbed Secondary Antibody, Alexa Fluor 647 (ThermoFisher, A-21236)

- Goat anti-Rabbit IgG (H+L) Highly Cross-Adsorbed Secondary Antibody, Alexa Fluor 488 (ThermoFisher, A-11034)

- Goat anti-Rabbit IgG (H+L) Highly Cross-Adsorbed Secondary Antibody, Alexa Fluor 647 (ThermoFisher, A-21245)

- Donkey anti-Rabbit IgG (H+L) Highly Cross-Adsorbed Secondary Antibody, Alexa Fluor 568 (ThermoFisher, A10042)

- Donkey anti-Rat IgG (H+L) Highly Cross-Adsorbed Secondary Antibody, Alexa Fluor 488 (ThermoFisher, A-21208)

- DAPI and Hoechst Nucleic Acid Stains (ThermoFisher, H3569)

### Live imaging of organoids

Bright-field images of live cerebral organoids were acquired with an epifluorescence microscope using a 10x air objective. The focus was manually adjusted.

### Live imaging of cell divisions in organoids

To observe cleavage planes in dividing cells, cerebral organoids were incubated with 1 μM SiR-tubulin (SpiroChrome, sc002) for 1 h before imaging, or with 0.5 μg/ml Hoechst 33342 (ThermoFisher, 62249) and 1 μM SiR-tubulin. Live organoids were imaged using the Olympus FV3000 confocal microscope with the UPLSAPO30XS (silicon oil) objective. Organoids were imaged for 2-8 h at 37 °C and 5% CO_2_.

### Image analysis

#### Organoid 3D morphometry

3D segmentation was performed using a custom pipeline with 3D stacks acquired at 2 μm/pixel and 4 μm/pixel z-resolution. Images acquired using the Luxendo MuViSPIM microscope were captured from two angles, and both views were merged into one before segmentation. When necessary, the image tiles were stitched together using a pairwise stitching approach(*42*). The stacks were then rescaled to a 1:1:1 voxel ratio (2 μm/pixel or 4 μm/pixel) before segmentation. Segmentation was carried out using a U-Net algorithm(*43*). When necessary, the segmented images were reviewed and manually corrected to ensure segmentation accuracy. Segmented images were converted to triangular meshes, and geometric parameters were obtained by adapting the approach we previously described(*28*) into a Python-based pipeline that incorporated scikit-image(*44*) and trimesh v4.11.2(*45*). For all measurements, quality control of the segmentation was carried out by discarding lumens with a volume less than 0.5 nl.

The following geometric parameters were measured:

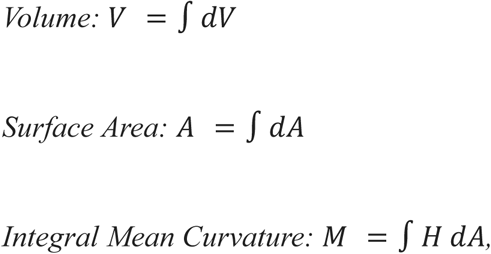

where H is the mean curvature.

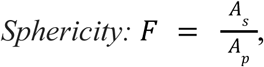

where As is the surface area of a sphere with the same volume as the object, and Ap is the measured surface area.

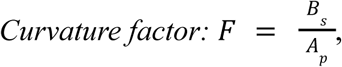

where Bs is the surface area of a sphere with the same integral mean curvature as the object, and Ap is the measured surface area.

Weighted mean by surface area parameters were calculated as follows:

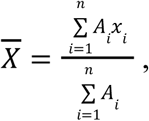

where x is the measured value, A_i_ is the measured surface area (weight), n is the number of objects.

#### Organoid 2D analysis

For quantifying neural bud circularity, neural bud thickness and the coefficient of variation of neural buds thickness, the z-plane containing the largest cross-sectional area of each bud within the organoid was selected. Neural bud thickness and the coefficient of variation were measured using custom codes developed in Fiji(*46*) and Python. The circularity of the buds was calculated from the thickness image masks:

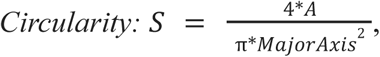

where *MajorAxis* is the longest diameter of an ellipse fitted to the object.

Organoid area measurements were performed using custom Python code. After segmentation, each image was manually checked for accuracy.

In experiments involving OptoShroom3 organoids, lumen perimeter measurements were performed using Fiji, with the GFP-N-Shroom3-iLID channel for the ON condition and the SiR-tubulin channel for the Control condition.

#### Organoid cell fate and division mode analysis

The numbers of PH3- and TBR2-positive cells were counted manually using Fiji. To quantify abventricular dividing cells in each neural bud of brain organoid, the z-plane with the largest cross-sectional area for each bud was selected. The orientation of cleavage planes in dividing cells was measured based on the SiR-tubulin signal during anaphase or telophase. The lumen surface was used as the reference surface for these measurements. The coordinates of two vectors were obtained from the images: one vector represented the cleavage plane, and the other corresponded to the lumen surface. The angle between them was then calculated.

### Quantitative RT-PCR

Total RNA was extracted from 30 cerebral organoids per condition in each experiment using the RNeasy Micro Kit (QIAGEN, 74004) according to the manufacturer’s instructions. RNA concentrations ranged from 600 to 1000 ng/μl. cDNA was synthesized from 1 μg of total RNA using the PrimeScript 1st strand cDNA Synthesis Kit (TaKaRa, 6110A). Quantitative RT-PCR was performed using LightCycler 480 SYBR Green I Master (Roche, 4707516001) with gene-specific primers on a LightCycler 480 System (Roche). The expression levels of the target genes were normalized to GAPDH.

*Primers for RT-PCR:*

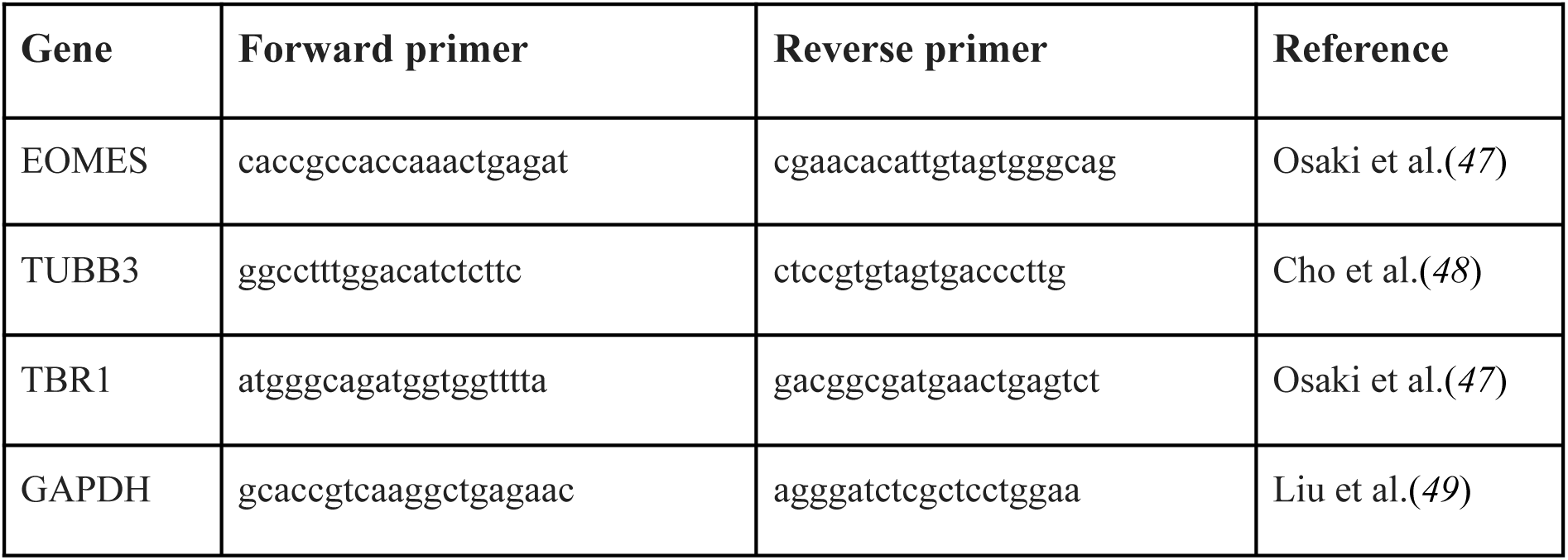

### Statistics and reproducibility

No statistical method was used to predetermine sample size. Statistical analyses were performed using the Mann-Whitney U test or two-sided paired t-test, as appropriate. Angular data were analyzed using the Watson-Williams test (a circular analogue of one-way ANOVA), after verifying the assumption of common concentration using the Wallraff test. A p-value < 0.05 was considered statistically significant. All experiments were reproducible, and the sample size is indicated in the figure legends.

Image intensity thresholds were adjusted for figure presentation. These adjustments were used for display only and not for quantitative analysis.

## Generative AI and AI-assisted technologies

AI tools were used to improve text clarity and assist with code generation. The authors reviewed and edited all outputs and assume full responsibility for the content.

## Acknowledgments

We are grateful to M. Matsuda and M. Sanaki for their experimental help; to M. Albert and M. Lancaster for their advice on organoid culture and analysis; to L. Rappez for his advice on lumen segmentation; to MPI-CBG Organoid and Stem Cell Facility for cell culture support; to MPI-CBG Light Microscopy, EMBL Barcelona Mesoscopic Imaging Facility, and PoL Microscopy Facility for imaging support. This work was supported by the Deutsche Forschungsgemeinschaft (DFG, German Research Foundation) under Germany’s Excellence Strategy - EXC 2068 - 390729961 - Cluster of Excellence Physics of Life of TU Dresden; the European Research Council (ERC) under the European Union’s Horizon 2020 research and innovation program (grant agreement No. 101002564 to M.E.); American Heart Association award (25CDA1432232 to K.I.). M.E. is supported by the Alexander von Humboldt Foundation in the framework of the Alexander von Humboldt Professorship endowed by the Federal Ministry of Education and Research. M.M. is supported by the Dresden International Graduate School for Interdisciplinary Life Sciences (DIGS-ILS).

## Author contributions

M.M. performed most of the experiments and analyses. G.M-A. helped with cell cloning, analysis and segmentation pipeline. K.I. helped with analysis and segmentation pipeline. J.P. helped with image analysis. M.E. supervised the project. All authors contributed to the manuscript and approved the final version.

## Data and materials availability

The custom scripts used are available from Github [https://github.com/mebisuya/BrainOrganoidShape].

## Supplementary figure legends

**Supplementary Fig. S1 Generation and characterization of human cerebral organoids with inducible Shroom3 stabilization**

(A) Protocol for generating cerebral organoids from human iPSCs. Colors indicate changes in media and developmental stages (see Methods). Organoids were embedded in 100% Matrigel on day 7, and apical constriction was induced on day 8 in the Shroom3 ON condition by adding 10 μM TMP. Scale bar: 100 μm. (B) Representative immunofluorescence images of cryosections of neural buds on day 15, labeled for TUJ1 (early-born neurons, magenta), PAX6 (neural progenitor cells, gray), and ZO-1 (apical marker, green). Yellow arrows indicate areas with TUJ1-positive cells. Scale bar: 100 μm. (C) Representative immunofluorescence images of iPSCs expressing Shroom3-V5-DHFR construct. The iPS cells were treated with TMP for 24 h and stained for ZO-1 (apical marker, green) and V5-tag (Shroom3, gray). The Scale bar: 20 μm. (D) Representative light-sheet immunofluorescence images of day 9 organoids, labeled for ZO-1 and V5-tag. Scale bar: 100 μm.

**Supplementary Fig. S2 Computational segmentation and morphometric analysis**

(A) Segmentation pipeline for cerebral organoids based on U-net segmentation and mesh approach. (B) Quantification of lumen geometric parameters in day 10 and 15 organoids. N = 94-186, 11-26 organoids per condition. Part of day 15 data are also shown in Fig. 1F. (C) Quantification of thickness of neural buds across 3 days. N = 48-172, 9-13 organoids per condition. (D) Quantification of organoid geometric parameters in days 10 and 15 organoids. N = 11-26. (E) Representative immunofluorescence light-sheet images of day 12 organoids from another Shroom3-DHFR clonal line (Clone 2) (left). Quantification of circularity of neural buds (right). N = 19-38, 5-9 organoids per condition. Scale bar: 100 μm. (F) Representative immunofluorescence light-sheet images of day 15 organoids from wild-type (WT) cells in the absence and presence of TMP (left). Quantification of circularity of neural buds (right). N = 26-39, 7-10 organoids per condition. Scale bar: 100 μm. For all graphs, the median and IQR are displayed, and all statistical comparisons were performed using Mann-Whitney U test.

**Supplementary Fig. S3 Evaluation of apoptosis and intermediate progenitor abundance**

(A) Representative immunofluorescence images of neural bud cryosections labeled for cleaved CASPASE-3 (apoptosis marker, yellow) and ZO-1 (apical marker, green). Blue arrows indicate apoptotic cells. Scale bar: 100 μm. (B) Representative immunofluorescence images of neural bud cryosections, labeled for TBR2 (intermediate progenitors, yellow) and DAPI (nuclei, blue) in day 15 organoids from another Shroom3-DHFR clonal line (Clone 2) (left). White arrows indicate TBR2-positive cells. Scale bar: 100 μm. Quantification of TBR2-positive cells normalized to bud area (right). N = 19-25, 10-14 organoids per condition. (C) Representative immunofluorescence images of neural bud cryosection in day 15 organoids from WT cells in the absence and presence of TMP (left). White arrows indicate TBR2-positive cells. Scale bar: 100 μm. Quantification of TBR2-positive cells normalized to bud area (right). N = 91-98, 14-16 organoids per condition. For all graphs, the median and IQR are displayed, and all statistical comparisons were performed using Mann-Whitney U test.

**Supplementary Fig. S4 Cleavage plane orientation analysis**

Distribution of cleavage plane orientations in Control and Shroom3 ON conditions. Measurements were taken at the anaphase, when microtubules start to separate. Angles near 90° correspond to vertical cleavage planes, whereas angles near 0° correspond to horizontal cleavage planes. Angles were grouped into seven categories: 0°, 15°, 30°, 45°, 60°, 75°, and 90°. N = 3-4 organoids per condition per experiment. The total sum of data is shown in Fig. 3B.

**Supplementary Fig. S5 Optogenetic activation of Shroom3 and cleavage plane orientation analysis**

(A) Representative live-imaging frames of non-illuminated (Control) and illuminated (ON) neural buds within the same OptoShroom3 organoid on day 12. Mitotic spindles were visualized with SiR-tubulin, and green dashed lines indicate the lumen surface. Light stimulation was applied in 9-min ON / 1-min OFF cycles. The left and right images show the start and end of 60 min of photostimulation, respectively. The illuminated neural bud (ON) is the same sample shown in Fig. 3D. Scale bar: 10 μm. (B) Distribution of cleavage plane orientations in non-illuminated (Control) and illuminated (ON) buds within the same OptoShroom3 organoids. Measurements were taken at the anaphase, when microtubules start to separate. Angles near 90° correspond to vertical cleavage planes, whereas angles near 0° correspond to horizontal cleavage planes. Angles were grouped into seven categories: 0°, 15°, 30°, 45°, 60°, 75°, and 90°. N = 9 neural buds per condition. The total sum of data is shown in Fig. 3F.

**Movie S1**

3D reconstructions of lumens in Control and Shroom3 ON organoids on day 10.

**Movie S2**

Live imaging of Control and Shroom3 ON organoids on day 12. White arrows indicate mitotic spindles, with cleavage plane angles shown and categorised into seven groups: 0°, 15°, 30°, 45°, 60°, 75° and 90°. Organoids were incubated with 1 μM SiR-tubulin before imaging. Snapshots of videos on the left are shown in Fig. 3A. Scale bar: 10 μm.

**Movie S3**

Live imaging of day 12 and day 14 OptoShroom3 neural buds during 60 min of photostimulation. Lumens are visualized with GFP-N-Shroom3-iLID. Snapshots of videos on the left are shown in Fig. 3D. Scale bar: 10 μm.

**Movie S4**

Live imaging of day 14 and day 15 OptoShroom3 neural buds after 1 h photostimulation. During photostimulation, buds within the same organoid were either non-illuminated (Control) or illuminated (ON). White arrows indicate mitotic spindles, with cleavage plane angles shown and categorised into seven groups: 0°, 15°, 30°, 45°, 60°, 75° and 90°. Organoids were incubated with 1 μM SiR-tubulin before imaging. The neural bud on the left (ON) is the same sample shown in movie S3. Scale bar: 10 μm.

