## Supplementary Figures S1-5 for "Synthetic lumen rounding directs neural progenitor division mode"

**Supplementary Figure 1**

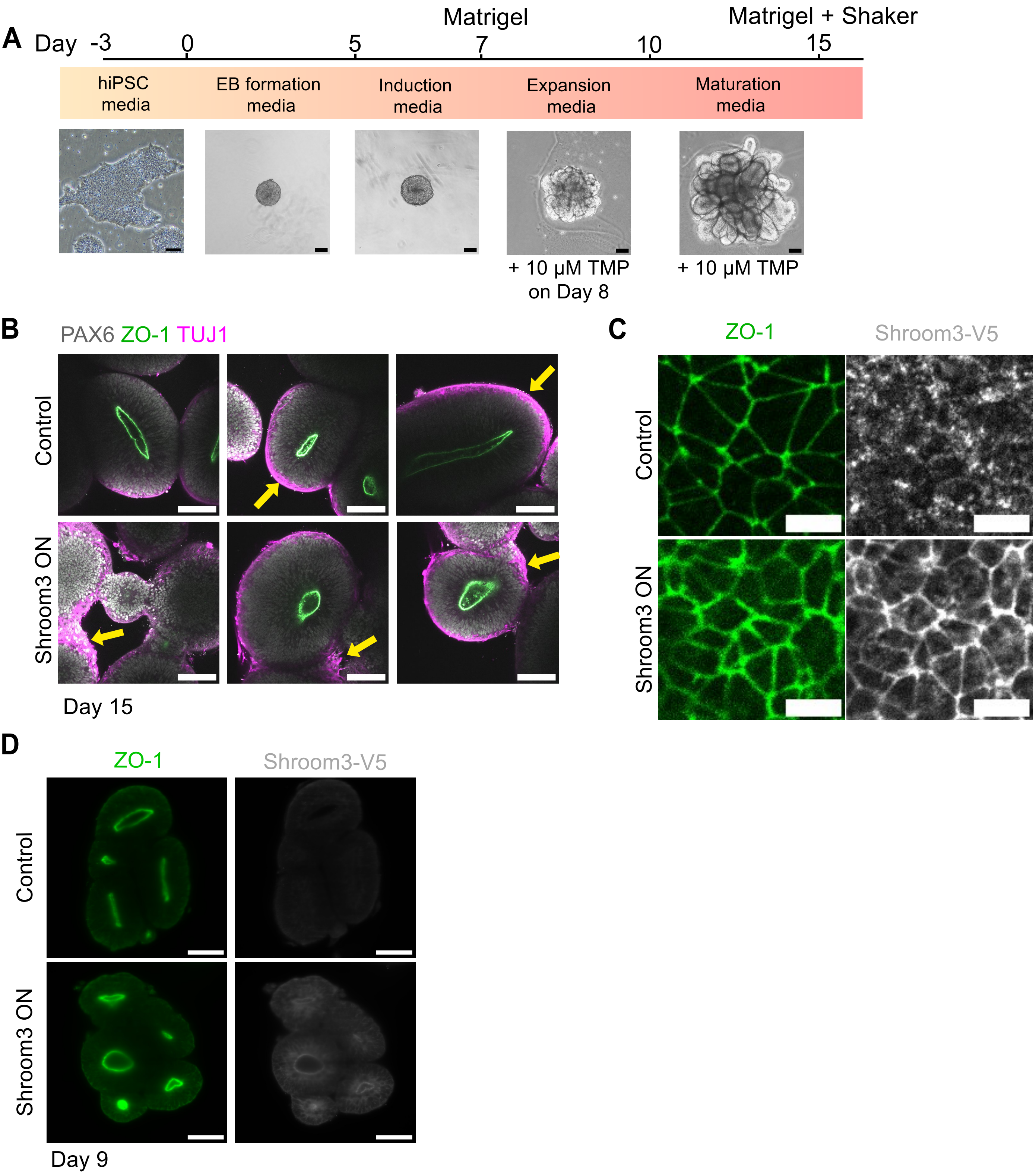

### Supplementary Figure 2

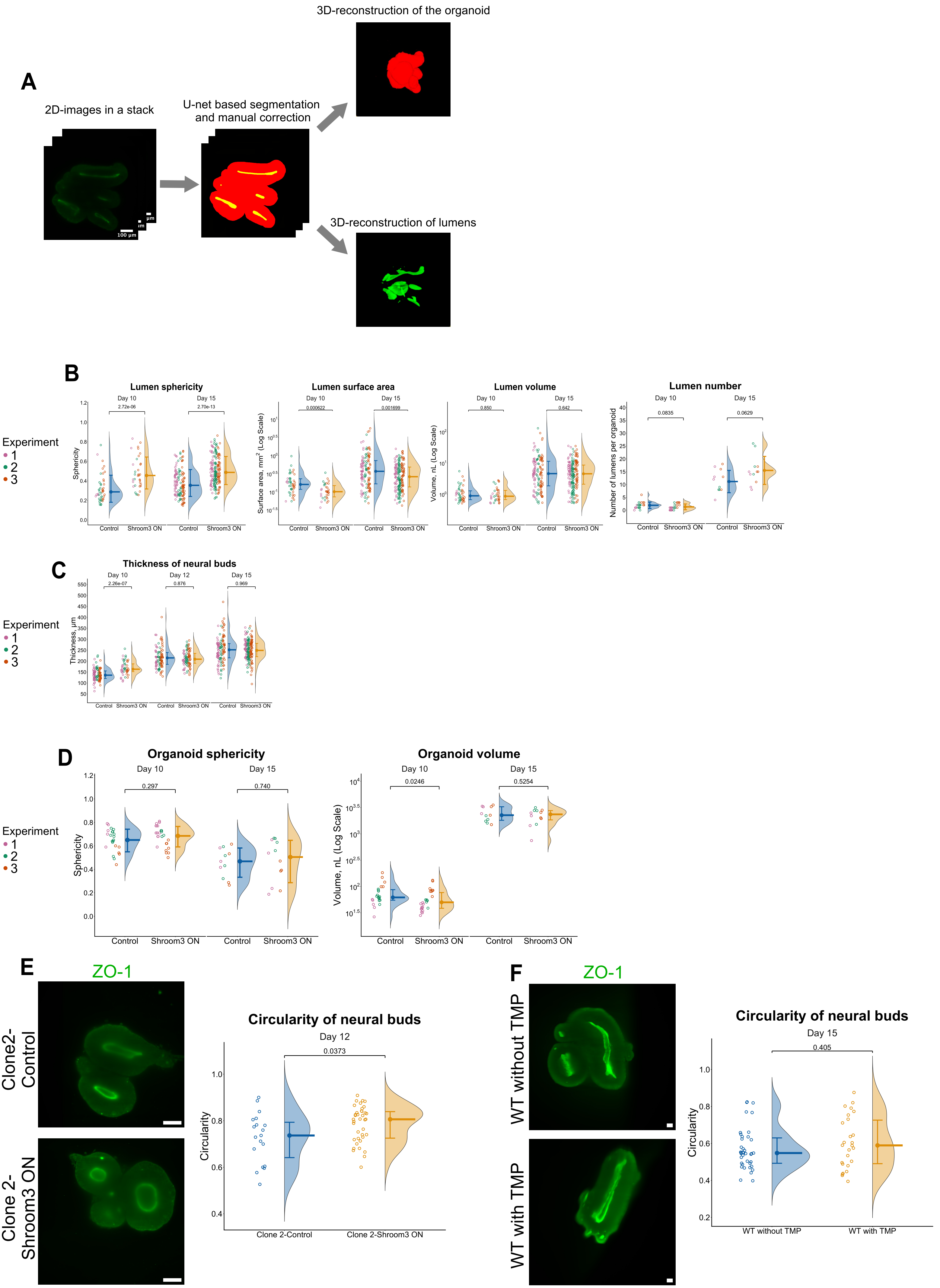

**Supplementary Figure 3**

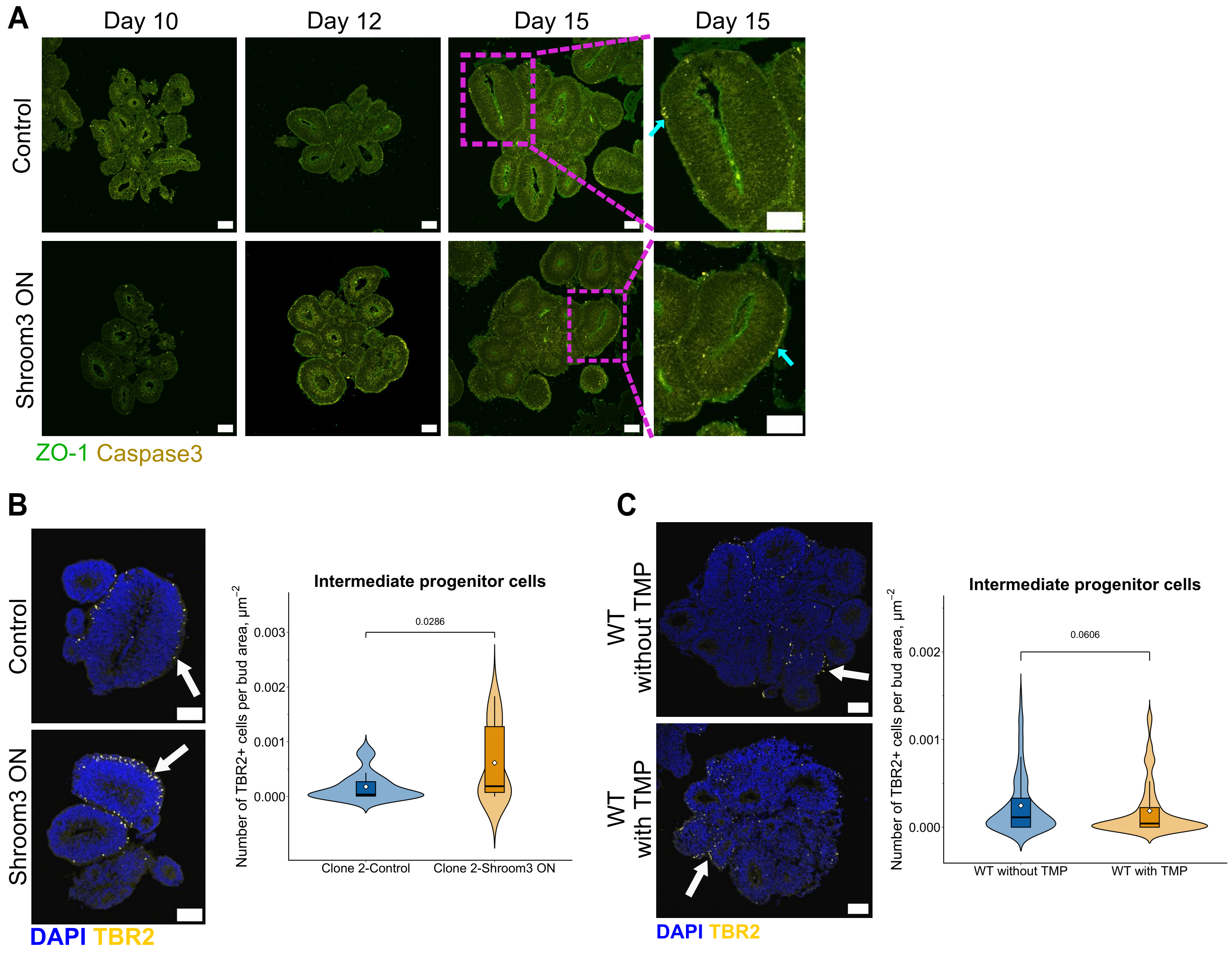

**Supplementary Figure 4**

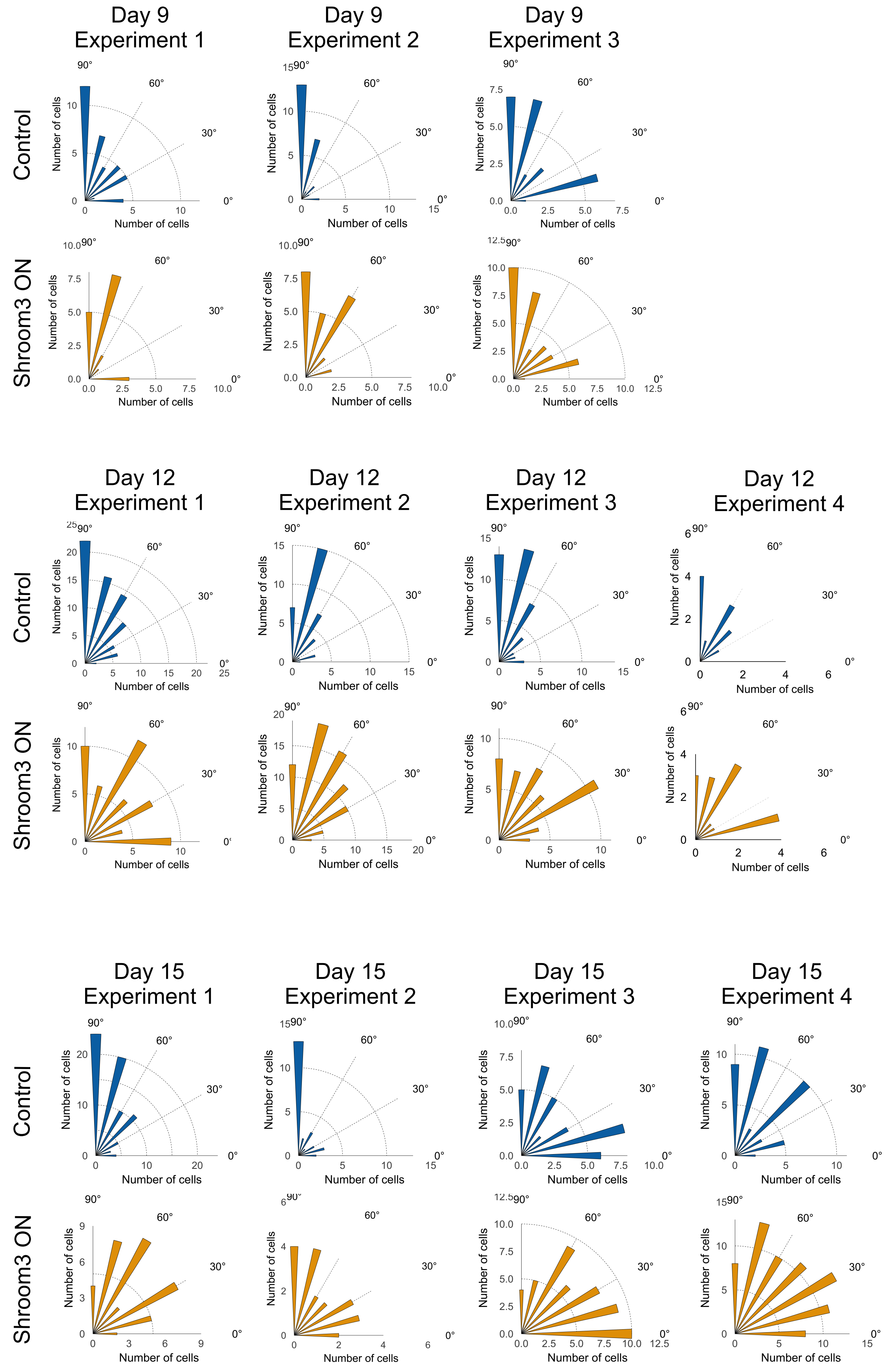

**Supplementary Figure 5**

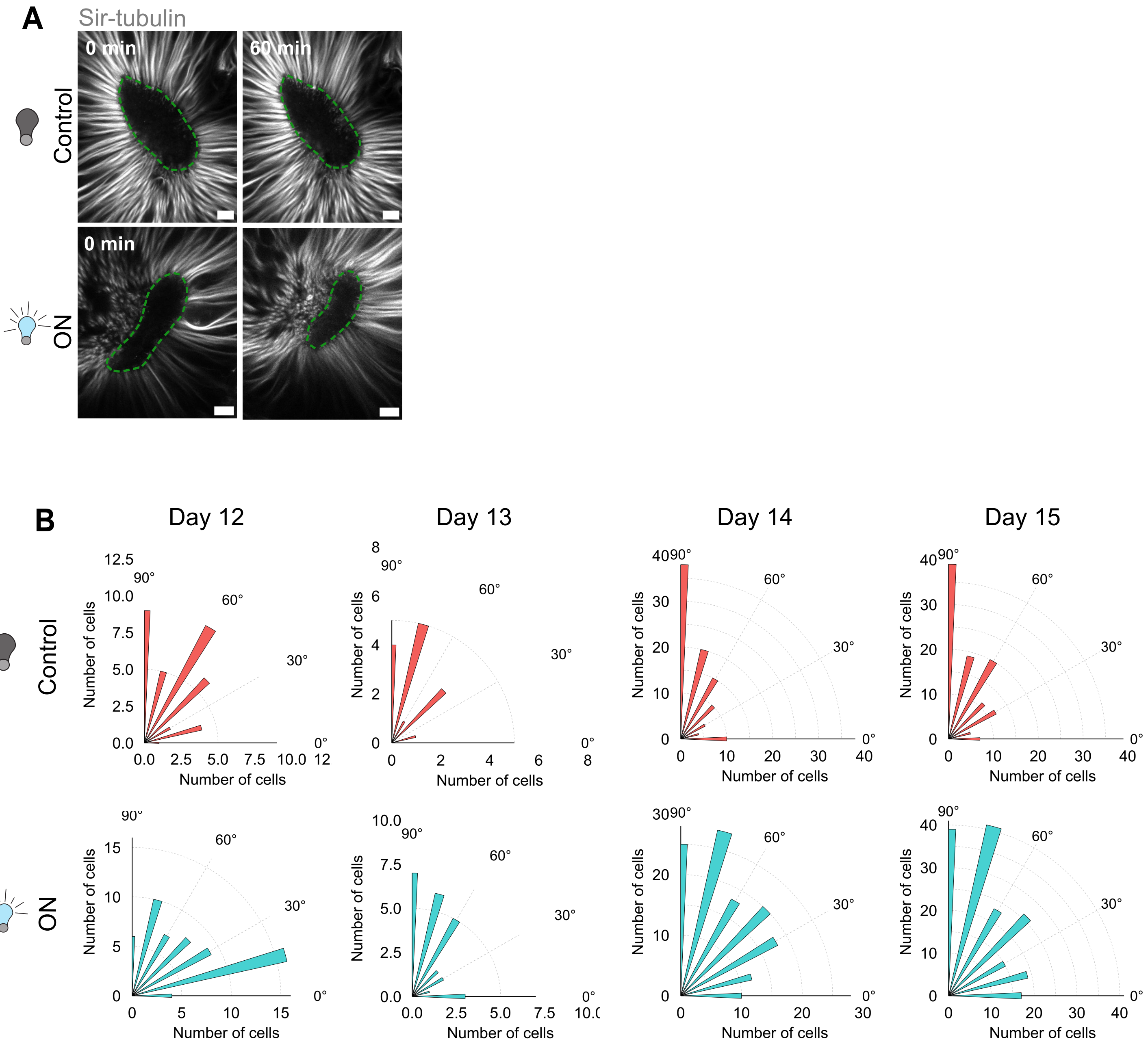
